# Catch me if you can: wild *Morpho* butterflies trade speed for erraticity in escape flight

**DOI:** 10.64898/2026.06.03.729813

**Authors:** Raphaël Dupillier, Violaine Llaurens, Florian Muijres, Vincent Debat

## Abstract

Predator–prey interactions shape the evolution of escape behavior in prey, including different combinations of evasive movements, that may enhance unpredictability in fleeing directions and trajectories. So-called protean motion can enhance survival of flying prey in the wild, but quantifying such behaviors under natural conditions remains challenging. Here we used stereoscopic high-speed videography to record the escape flight behavior of wild males of the butterfly species *Morpho menelaus* in the Amazonian rainforest, and reconstructed 3D flight trajectories using artificial-neural-network-based tracking. During the experiments, we used a lure to attract freely patrolling male butterflies and elicited escape flights by intercepting their trajectory with a looming insect net swing. We then compared the escape flight kinematics to the pre-attack patrolling behavior. Attacks first induced a rapid upward maneuvering, directly followed by an unpredictable horizontal turn. The following escape flight trajectories showed increased horizontal erraticity and greater intra-individual heading variability, as compared to the pre-attack flight. Surprisingly, the mean speed decreased in the escape phase, notably in the horizontal plane. A significant negative association between horizontal trajectory complexity and flight speed was detected, indicating a speed-erraticity trade-off. These results show that wild *Morpho* butterflies respond to attacks by combining a climbing maneuver with an unpredictable heading change, followed by a protean escape flight; this increased escape erraticity comes at the expense of reduced escape flight speed. Because these large and relatively slow-flying butterflies display bright iridescent blue coloration on their dorsal wing side, erraticity during flight might enhance the dynamic flash coloration, likely limiting accurate targeting by predators.

## Introduction

Predator-prey interactions impose strong selective pressures shaping the evolution of multiple behavioral and morphological traits in animals (Schmitz, 2017). In particular, traits enhancing evasive success are likely under strong selection in prey species. Studying evasive behaviors in prey thus allows estimating the effect of predator selection on the evolution of traits enhancing survival in the wild, such as acceleration capacity or unpredictability in evasive maneuvers and escape trajectories.

Movement unpredictability has been documented to increase escape success in many animals (Humphries & Driver, 1970). Because predators often rely on their ability to accurately predict the position of their moving prey to perform efficient strikes, unpredictable prey behaviors can swindle the aiming system of predators (Combes et al., 2013). Such unpredictability might be especially efficient when facing predators using ballistic interception (Moore & Biewener, 2015), *i.e.* hunting behaviors relying on the estimation of an intercepting trajectory before launching a rapid bursting attack. Furthermore, because predators can learn from previously failed attacks, unpredictable escapes can limit attack success performed by specialized predators (Edut & Eilam, 2004).

Unpredictability in evasive behaviors of prey can combine different traits enhancing (1) directional variability when performing an evasive maneuver, and (2) protean (i.e. erratic) motion during the subsequent escape.

1. Once a prey becomes aware of an incoming threat, it often first generates a rapid evasive maneuver (Domenici et al., 2011b). Unpredictability of this initial response has been shown to be a common and important factor in increasing escape success (e.g. Domenici et al., 2008). Indeed, prey escaping in highly varying directions after each attack makes it harder for predators to anticipate the future position of the prey. However, in a review on evasive behavior directions, Domenici et al. (2011a) highlighted some costs associated with unpredictability in the direction of escape trajectories: in species where individuals escape in completely random directions, some individuals will eventually flee towards the threat rather than away from it (*e.g.* in the larvae of the mosquito *Culex pipiens*, Brackenbury, 1999). The resulting escape model proposed in Domenici et al. (2011a) thus indicates that escape success is maximal when prey flee in unpredictable directions biased away from the threat. This model aligns well with empirical data, including escaping lizards (Cooper & Sherbrooke, 2016), fish (Domenici & Hale, 2019) and grasshoppers (Bateman & Fleming, 2014), but with its limitations (Corcoran & Conner, 2016). A critical aspect that interferes with this model is the initial direction and speed of moving prey that may limit the span of escape angles due to inertia (Wynn et al., 2015).
2. After this initial evasive maneuver, escaping animals tend to perform complex high-speed escape movements, often with unpredictable trajectories. Both the increase in speed and erraticity can contribute to evasive success (Clemente & Wilson, 2016). However, increasing speed may be limited by evolutionary or immediate constraints, including morphological (Koehl, 1996), physiological or environmental conditions (Crall et al., 2015). In such cases, protean movements may enhance escape performance more effectively than fast movements (A. M. Wilson et al., 2018). Because rapid turning imposes force and work that rise with speed (Alexander, 1982), prey cannot maximize both speed and erraticity simultaneously; instead, they tend to operate along a speed–maneuverability continuum tuned by ecology and morphology (e.g. Clemente & Wilson, 2015; Moore et al., 2017; Mountcastle et al., 2015). In butterflies with low wing loading, selection may therefore favor enhanced erraticity over peak flight speed when escaping from interception-based predators.

Moreover, during escape, travelling in a straight line at a constant speed is also easier for predators to detect and track (Brunyé et al., 2019), whereas prey that display greater erraticity in their movements tend to disorient predators and reduce their strike accuracy (Richardson et al., 2018). Several insect prey species indeed display baseline protean movements enhancing their evasive success, including fruit flies (Combes et al., 2012) and mosquitoes (Cribellier et al., 2022). The evolution of faster versus maneuverable escape locomotion in different species therefore depends on the ecological context encountered, but also on the baseline level of erraticity performed.

Among flying animals, butterflies typically display highly maneuverable flap-gliding flight, and their escape is often described as “erratic” (Chai & Srygley, 1990). This maneuverability likely arises from their distinctive morphology, characterized by large wings and a relatively small body, hence a low wing loading (Le Roy et al., 2019). Nevertheless, the levels of unpredictability in their trajectories in regular versus stressful conditions have never been explicitly characterized.

Here we focus on *Morpho menelaus*, a large butterfly species of the *Morpho* genus with a low wing-loading. These butterflies live in the understory of Neotropical rainforests of central and south America, where their main predators are thought to be perching birds that rely on ballistic interception, such as jacamars and kingbirds (Chai, 1986; Pinheiro, 1996). In *Morpho menelaus,* flight has been characterized in cage conditions where a fast, maneuverable flapping flight behavior has been documented (Le Roy, Amadori, et al., 2021). However, the change in flight behavior during predator attacks and the properties of its escape flight have never been explored. Given their large wings and low wing loading, we expect that *Morphos* will resolve the speed–erraticity tradeoff during escape flights by prioritizing erratic, protean behavior over flight speed. This prediction follows from the mechanical cost of turning at high speed and empirical demonstrations of speed–maneuverability compromises in other taxa (Clemente & Wilson, 2015; Moore & Biewener, 2015; Wynn et al., 2015).

Furthermore, *Morpho menelaus* butterflies typically fly in the understory strata of the forest, so evasive turning maneuvers might be biased towards more cluttered spaces that can serve as shelters (Wirsing et al., 2010). The dense vegetation encountered in the understory may thus also promote maneuverability (Crall et al., 2015), and result in erratic escaping movements rather than faster and straighter escape trajectories.

Erraticity in flight behavior might also be particularly advantageous in this species exhibiting iridescent blue coloration on the dorsal surface of the wings combined with brown ventral patterns. This strong dorsal-ventral color contrast generates conspicuous flashes as the butterfly beats its wings, and the combination of dynamic flashes and protean movements likely diminishes the targeting efficiency of predators (Young 1971; Debat et al., 2018). Both the understory habitat and the co-evolution with wing coloration makes the comparison of baseline *versus* escape flight unpredictability specifically relevant in *Morpho menelaus*.

Together, these considerations lead to a specific prediction: in *Morpho menelaus*, escape trajectories should become more erratic at the expense of a limited increase in flight speed, reflecting a speed–erraticity trade-off under interception risk.

Here, we tested this prediction by quantifying the unpredictability in flight behavior in wild *Morpho menelaus* butterflies when they are threatened, and how this correlates with escape flight speed. We recorded their flight in natural conditions to avoid spatial limitations that would hinder flight and are documented to bias the natural behavior of wild animals (Morgan & Tromborg, 2007). We specifically tested three hypotheses about the escape response of *Morpho* butterflies:

1. Escape flight unpredictability We expect *Morpho menelaus* to increase their escape unpredictability using two complementary mechanisms:
  a. We expect butterflies to initiate their escape using a rapid evasive turn in a stochastic direction, with a bias toward the visually cluttered forest understory, reducing predator tracking efficiency (Domenici & Ruxton, 2015).
  b. We expect an increase in flight path erraticity during the escape flights following the initial evasive maneuver, because higher path complexity decreases interception success (Richardson et al., 2018).
2. Escape flight speed We expect butterflies to increase their escape flight speed in response to a perceived threat, reducing capture probability by avian predators (Clemente & Wilson, 2015).
3. Trade–of f between escape flight speed and erraticity. We hypothesize a trade–off between increased escape flight speed and erraticity (Clemente & Wilson, 2015). For *Morpho menelaus*, we expect this trade-off to shift toward enhanced erraticity because the combination between wingbeat-induced blue flashing and trajectory erraticity amplifies visual signal disruption in predators (Murali & Kodandaramaiah, 2020).

We tested these three hypotheses by tracking multiple escape flights of *Morpho menelaus* in the wild. From these trajectories we quantified the changes in flight direction during evasive turns, and measured flight path erraticity, flight speed, and aerodynamic force production across the subsequent escape maneuvers.

## Methods

### 1. Experimental animals & conditions

All escape flights of *Morpho menelaus* butterflies were recorded in the Amazonian rainforest at the same locality on the Kaw mountain in French Guyana (4°32’41“N 52°09’42”W; Fig. 1a). Filming was performed on a partially dried-up riverbed, where *Morpho* males regularly patrol (Bouinier et al., 2025) (Fig. 1A). Experiments were performed during the two-month period of October and November 2024, in predominantly sunny conditions because males usually do not perform patrolling flight in cloudy or rainy conditions. The filming hours correspond to the natural patrolling hours of *M. menelaus* (Bouinier et al., 2025), i.e. between 9 to 11 am.

**Figure 1:**
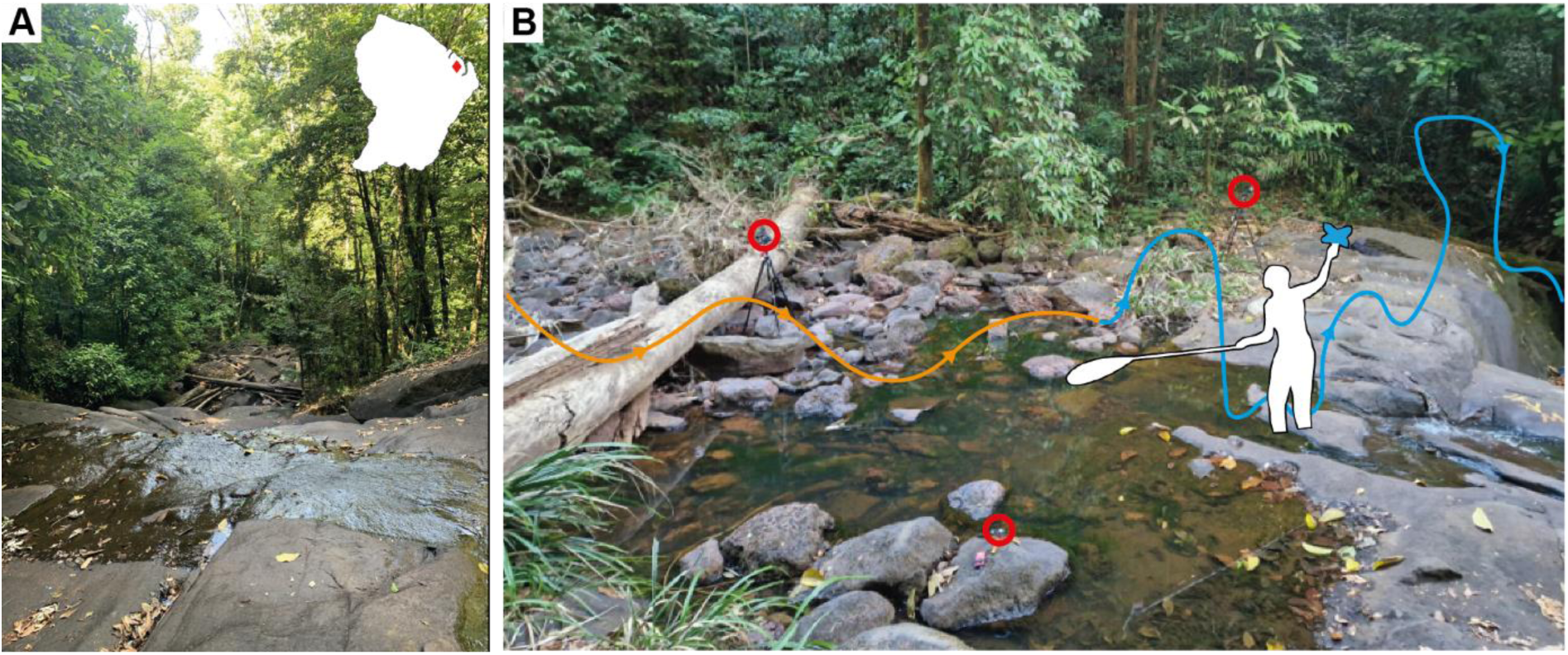
Filming location and experimental set-up. All *Morpho* butterflies were filmed at this riverbed, among the dense Amazonian rainforest in the Kaw mountain of French Guiana. **A)** View of the axis of the river in the middle of the filming set-up, illustrating the natural corridor along which Morpho butterfly males regularly patrol and were spotted for our experiment. On the top right, we show the map of French Guiana, with the location of this riverbed indicated by the red diamond (*4°32’41“N 52°09’42”W*). **B)** Image of the stereoscopic videography set-up in the wild with a representation of a typical *Morpho* flight sequence. The positions of the three GoPro cameras are indicated by the red circles with their respective orientations aligned on the center of the filming space, represented by the outline of one of the experimenters. A typical flight sequence proceeds as follows: An experimenter would be placed in the center of the filmed space, waving a blue shiny lure in the direction of an incoming *Morpho*. Attracted to the lure, the butterfly enters the filming space with an undisturbed flight illustrated by the orange trajectory. Once close enough, the experimenter simulates an attack by waving a tied-up entomological net in the direction of the butterfly, triggering a rapid evasive maneuver and a subsequent escape flight illustrated by the blue trajectory.

Although we only recorded one escape flight per sequence, we cannot fully discard pseudo-replication as we could not identify the different wild individuals: the same male could have been filmed on different days. Previous mark-recapture data showed that *M. menelaus* had a rate of recapture along consecutive days of about 0.17 (Le Roy, Roux, et al., 2021).

### 2. Experimental setup & procedure

#### Experimental setup

Our experimental setup consisted of a high-speed videography system placed across a riverbed along which *Morpho* males regularly patrol (Fig. 1b). The videography system consisted of three Gopro cameras (one Hero8 black, two Hero11 black), all recording at a spatial resolution of 2704 x 1520 pixels and a temporal resolution of 120 frames per second. The cameras were positioned to maximize the diversity of view angles of a filming space of about 7 m^3^.

#### Experimental procedure

At the start of each experimental day, we installed the experimental setup, performed our video calibration routine (see below), started our experimental video recordings at 8:30 am, and then waited for patrolling butterflies to pass by.

When a butterfly appeared, we would first wave the blue iridescent lure to attract the butterfly within the field of view of the three cameras. Once a butterfly was near, we would simulate a predator attack by swiftly waving a tied-up entomological net towards the animal. The net would generally come from below, and would trigger a distinct rapid evasive response, and a subsequent escape flight. As such, each movie captured an initial flight directed towards the lure followed by a disturbed, rapid evasive flight maneuver, and finally the subsequent escape flight. After a successful recording, the experimenter would operate a clapper board in the middle of the filming region for video frame synchronization. Because Gopro cameras do not have the option for multi-camera video frame synchronization, the clapperboard instances allow for aligning the videos captured by each of the three cameras into a single sequence.

### 3. Calibration of the camera system

The stereoscopic camera system was calibrated using a dedicated calibration procedure. First, the cameras’ lens distortions were corrected using GoPro’s official Plug-in for Adobe Premiere. Second, we used a series of calibration videos to perform the three-dimensional calibration, as described in Theriault et al. (2014).

Before and after each daily experimental session, we recorded dedicated calibration videos for that day. For this, we moved a 0.9m T-shaped calibration wand attached to a long rod (> 4m) across the total filming volume and dropped an object in the middle of this volume. On each video, we then automatically tracked the position of both wand extremities using the deep learning open-source python toolbox DeepLabCut (DLC, Mathis et al., 2018; Nath et al., 2019).

Because the complexity of the environment caused very different backgrounds for each camera angle, we trained a distinct tracking model for each camera view. The training was performed on 100 frames picked randomly from all the calibration sequences. From the three sets of 2D positions of the wand extremities obtained by DLC, we retrieved the extrinsic and intrinsic parameters of the cameras in a transformation matrix through the direct linear transformation process using the EasyWand MATLAB application (Theriault et al., 2014). The precision of our 3D digitalization was estimated to have a mean error of 4.5 mm.

Finally, we defined the vertical axis in our world reference frame by tracking the object that we dropped during the calibration video recordings.

### 4. Flight trajectory reconstruction and analysis

To reconstruct the 3D flight trajectories, we first tracked the position of the butterflies on each camera view in 2D. As for the calibration, we used DeepLabCut to automatize the butterfly tracking, with a distinct model trained for each camera view. 150 manually-digitized video frames were used for the training, picked based on the visibility of the butterfly across all 32 recorded trajectories. The tracked positions were filtered following the base 0.6 likelihood cutoff value. The positions that did not reach this value were completed manually with the MATLAB application DLTdv8 (Hedrick, 2008). To shorten the manual tracking, only one out of five video frames was tracked manually, and the resulting small gaps were completed by spline interpolation. This method also allowed us to complete missing positions for when the butterfly became invisible in the frame (e.g. when passing behind an obstacle). However, because spline interpolation can affect variation compared to the real trajectory, a threshold of maximum 11 frames was set to limit the size of gaps over which interpolation could securely be performed. This value corresponds to half of the average wingbeat duration of *Morpho menelaus*.

Three-dimensional trajectories were reconstructed with the transformation matrix obtained via the calibration process. The reconstruction was done with the MATLAB application DLTdv8, producing the right-handed XYZ-coordinates of the butterfly at each frame. Here, X and Y define the horizontal plane, whereby the X-axis was approximately parallel to the riverbed; the Z-axis corresponds to the vertical upward axis.

To filter out the positional noise likely resulting from reconstruction imprecision, we passed the 3D coordinates of each trajectory through a Kalman filter following the procedure described in Muijres et al. (2014). The output contained the smoothed temporal dynamic of positions **X**=(*x,y,z*), velocity **U**=(*u,v,w*) and accelerations **A**=(*a_x_, a_y_, a_z_*) in three dimensions. These were all defined in the right-handed world reference frame, of which the coordinate system has the origin at the butterfly position at *t*=0s (moment of attack), the *x*-axis along the riverbed and the *z*-axis vertically up.

### 5. Flight kinematics analysis

Flight sequences generally consisted of three main phases: (1) the pre-attack approach flight towards the lure, (2) the rapid evasive maneuver in response to the net attack, and (3) finally the post-attack escape flight away from the attacker. We therefore separated each recorded flight track into these phases, based on the net attack timing. We defined time *t*=0 s as the moment when the Euclidean distance between the butterfly and the net was minimal. The evasive maneuver phase was then defined as a 200 ms time window around that moment (*t*=[-0.1 – 0.1] s); the pre-attack and post-attack phases time periods were thus defined as *T*_pre_<-0.1 s and *T*_post_>0.1 s, respectively.

Throughout each flight track, we estimated the following flight kinematics parameters: flight heading ϕ, ascend angle γ, flight velocity (**U**), flight path erraticity (*E*), and the net weight-normalized aerodynamic force (**F**/*mg*), where *mg* is body weight.

#### Flight speed and velocity

We characterized the escape flight performance using flight speed, whereby we assumed that an increase in escape flight speed enhances escape performance. Speed *U* was defined as the vector length of the flight velocity **U**=(*u,v,w*) and separated into the horizontal component *U*_hor_ and vertical component *U*_ver_.

#### Flight heading & ascend angle

Based on velocity vectors, we also estimated the flight directions throughout each flight track. This flight direction was separated into the flight heading within the horizontal plane, and the ascend angle within the vertical plane. Heading was thus defined as the angle relative to the *x*-axis along the riverbed, and calculated as ϕ=atan2(*u,v*). Ascend angle was similarly estimated as γ =atan2(*U*_ver_*,U*_hor_).

To analyze the turn dynamics during the evasive maneuver and consecutive escape flight, we isolated the initial direction change performed by the butterflies in response to the attack, which we defined as the turn angle.

Next to the heading in the world reference frame, we determined the heading in two additional reference frames

1. To quantify turn dynamics during the evasive maneuver, we secondly defined heading relative to the pre-attack approach flight heading, by aligning all approach flights through setting the mean approach heading to zero. This resulted in the aligned headings (ϕ_align_) dataset. In this reference frame, the variations in escape headings then equal the turn dynamics.
2. To quantify the variations in horizontal turn angle magnitude, we used a third heading dataset in which we mirrored all turning maneuvers to the left. The resulting mirrored dataset (ϕ_mirror_) thus consists of aligned approach flight headings, with only net right-handed escape turns.

#### Aerodynamic force production

All flight maneuvers depend on the production of aerodynamic forces by the flying butterflies. To assess these force dynamics, we determined the net weight-normalized aerodynamic forces as

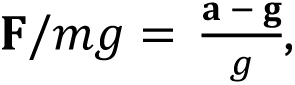

where *m* is body mass, **g** and *g* are the gravitational acceleration vector and scalar, respectively (**g**=(0,0,-*g*)), and **a** is the acceleration vector. From this normalized force vector, we determined the total force magnitude *F/mg* as the vector length, and the horizontal and vertical components as *F*_hor_/*mg* and *F*_ver_/*mg*, respectively.

#### Flight path erraticity

To estimate the erraticity of the trajectories, we measured a parameter based on positional-entropy following the derivation of Roberts et al. (2004) and Herbert-Read et al. (2017). Although this parameter was first applied to 2D data, its extension to 3D was implemented by Richardson et al. (2018). For this measurement, the successive positions of each butterfly were arranged into three embedding matrices, one for each dimension (𝑴_𝒙_, 𝑴_𝒚_, 𝑴_𝒛_) as illustrated below for the *x*-coordinates

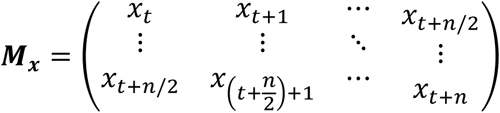

With *x_t_* the *x*-coordinate at time *t* and *n* the size of the trajectory over which erraticity is measured. The three embedding matrices are then horizontally concatenated into a single matrix **M**. For each column of **M** its mean was subtracted to focus on positional variations around the mean position during the time window corresponding to the associated column. The singular values of the resulting matrix **M**’ were then obtained by singular value decomposition (**UΣV\***). Entropy of the standardized singular values was then computed to capture the organization of successive three-dimensional positions, yielding a unique value used as erraticity index.

This measurement is highly dependent on the length of the trajectory. Because the different flight sequences had various durations, we measured the erraticity index within small segments of a fixed number of video frames along the full trajectory, advancing one frame at a time, effectively measuring the temporal dynamics of the trajectory’s erraticity. This method provides a quasi-continuous estimation of erraticity, therefore increasing the precision in comparison with the method presented by Herbert-Read et al. (2017). The size of the segment over which erraticity is measured must be long enough to allow the butterflies to perform erratic movements, as focusing on too-short segments may miss the actual complexity of the flight. We thus standardized the segment size to one-and-a-half wingbeat which corresponds to 33 video frames, assuming that butterflies generate erratic movements mainly through wing beating. Both horizontal and vertical components of erraticity were also measured distinctively by considering separately the (*x,y*) and the *z*-coordinates, respectively.

### 6. Statistical analysis

To explore how the escape flight of *Morpho menelaus* differs from its undisturbed approach flight, we performed statistical analyses focusing on the mean kinematics parameters during the pre-attack and post-attack phases.

All statistical testing was performed on Matlab v2024b, with the addition of the circular statistics toolbox where appropriate (Berens, 2009). For all comparative tests, we relied on both the *p*-value (*p*) and the effect size (*ES*). We follow the common guidelines for significance (*p*<0.05) and effect size interpretation: *ES*=0.1 to 0.3 for small effect sizes, *ES*=0.3 to 0.5 for medium effect sizes, and *ES*>0.5 for large effects. In the case of GLMs we also use the magnitude of the estimates of the model.

#### Escape heading & ascend angle

For our analysis of escape direction, because we separated horizontal and vertical directions, they were not measured in the same mathematical space. Horizontal directions were measured along a full circle making the measurement periodic; hence the horizontal angular data were treated using circular statistics. In contrast, vertical directions were bound between -90° and 90°, thus although they remain angular data, we analyzed them using linear statistics.

For the turn dynamics in both the horizontal and vertical plane, we performed the following tests:

- We tested whether the variability of flight angles changed between the pre-attack and post-attack phases. To test this, we measured the intra-individual standard deviation of angles (circular standard deviation for horizontal headings) of each phase and compared their distributions with a Wilcoxon signed-ranks test. The Wilcoxon signed-ranks test accounts for the low sample size of our dataset.

- We used a series of generalized linear models to test for the effect of both the approach direction and the attack direction on the escape direction (ascend angle and heading). Here, escape direction was set as the dependent parameter, and both approach direction and incoming attack direction were set as the independent parameters. For the vertical ascend angle, because they are bounded, we used a gaussian distribution for the model. For horizontal escape heading however, because they are circular hence periodic, we designed a GLM fitted for circular data based on the von Mises distribution (Fisher & Lee, 1992). In this case, the circular predictor values (i.e. approach direction & attack direction) were transformed into linear values by considering both their sine and cosine as described in Pewsey et al., (2013).

Specific for the turn dynamics in the horizontal plane, we performed two additional tests:

- To test for variations in turn dynamics within the horizontal plane, we used a circular V-test to test whether the aligned post-attack headings (ϕ_align,post_) differed from the initial 0° heading.

- To test whether butterflies escaped in a preferential direction in the surrounding environment we used the horizontal escape heading in the world reference frame (ϕ) and tested for uniformity using a Rayleigh’s test.

Specific for the turn dynamics in the vertical plane, we performed one additional test:

- To test for consistent ascend angle changes between the pre-attack and post-attack phases, we used a Wilcoxon signed ranks test on the mean ascend angle distributions.

#### Flight path erraticity

To determine whether erraticity increased after the attack, we compared the individual mean erraticity before and after the attack using a Wilcoxon signed ranks test. We performed these tests for the total erraticity, as well as the horizontal and vertical components separately.

#### Flight speed and aerodynamic forces

To investigate how force production and resulting flight speed varied during the escape, we compared the total, horizontal and vertical mean speed and aerodynamic force values between pre-attack and post-attack flight phases. Differences between phases were tested with a Wilcoxon signed-ranks test.

#### Testing for a tradeoff between escape flight unpredictability and speed

We hypothesize a trade–off between increased escape flight speed and flight path unpredictability in escaping *Morpho menelaus* (Clemente & Wilson, 2015). We tested for such trade-off by computing the correlation between post-attack flight path erraticity and escape flight speed using Spearman’s rank correlation coefficient.

Additionally, we tested the relationship between horizontal escape turn angle and flight speed before and after the attack, as larger turns could also correlate negatively with flight speed. This allowed us to assess whether incoming speed constrained escape direction and whether turn angle influenced subsequent flight speed. All correlations were performed using Spearman’s rank correlation coefficient.

Both the potential increase in escape flight speed and post-attack flight path erraticity should be powered by aerodynamic force production. Therefore, we used a pair of GLMs to test for the effect of aerodynamic force production on escape flight speed and erraticity during escape flight. In the first model, erraticity was set as the dependent parameters, and both aerodynamic force and speed as independent parameters (*E∼F/mg*+*U*). In the second model, we instead set speed as the dependent parameters, and both aerodynamic force and erraticity as independent parameters (*U∼F/mg*+*E*).

This approach allows us to evaluate the effect of force on each response variable while controlling for the other. Finally, we combined both models in a single visualization to show how the combination of speed and erraticity depend on aerodynamic force production, as *E+U*∼*F*/*mg*. With this multi-model architecture, we investigated the role of aerodynamic force production in modulating flight speed and flight path erraticity during the escape flights of *Morpho melenaus*. These tests were performed on total scalar values, and for the horizontal and vertical components, separately.

### 7. Use of AI

AI tools were used to support programming and manuscript preparation. During writing, AI was used for language editing and clarity only; all scientific content and interpretations were produced by the authors. During data analysis, AI-assisted tools were used to improve analysis code structure and efficiency, with all analytical decisions and validation performed by the authors.

## Results

We recorded 32 three-dimensional approach and escape flight sequences of wild *Morpho menelaus* males. The sequences comprised an approaching phase where butterflies headed toward the lure along a descending patrolling path, a rapid upward-directed evasive maneuver elicited by the looming net, followed by a subsequent escape flight phase (fig. 1b). We separated all tracks in the approach trajectories (*t* < 0 s), and post-attack escape flight trajectories (*t* > 0 s); *t* = 0 s is defined as the moment at which the distance between the butterfly and the net is the smallest. The resulting approach and escape trajectories lasted 0.57±0.26 s and 0.51±0.32 s, respectively (mean ± standard deviation).

### Ascend kinematics: Attacks trigger a rapid climbing maneuver

The overview of the vertical component of the filmed trajectories reveals that the recorded flights can be separated into four phases (fig.2A,B): (1) **descent** phase (*t*∼[-0.8; -0.3] s): the butterflies approached the lure using a descending flight path; (2) **evasive** maneuver phase (*t*∼[-0.3; 0.1]): the butterflies produced an increased upward-directed aerodynamic force, thereby changing the descending flight into a climbing maneuver. (3) **climbing** escape phase (*t*∼[0.1; 0.35] s): the butterflies rapidly climb during the initial part of the escape flight phase; (4) **horizontal** escape phase (*t*∼[0.35; 0.8] s): Finally, the escaping butterflies level off, as the ascend angle progressively stabilizes towards zero, resulting in a horizontal escape flight.

**Figure 2:**
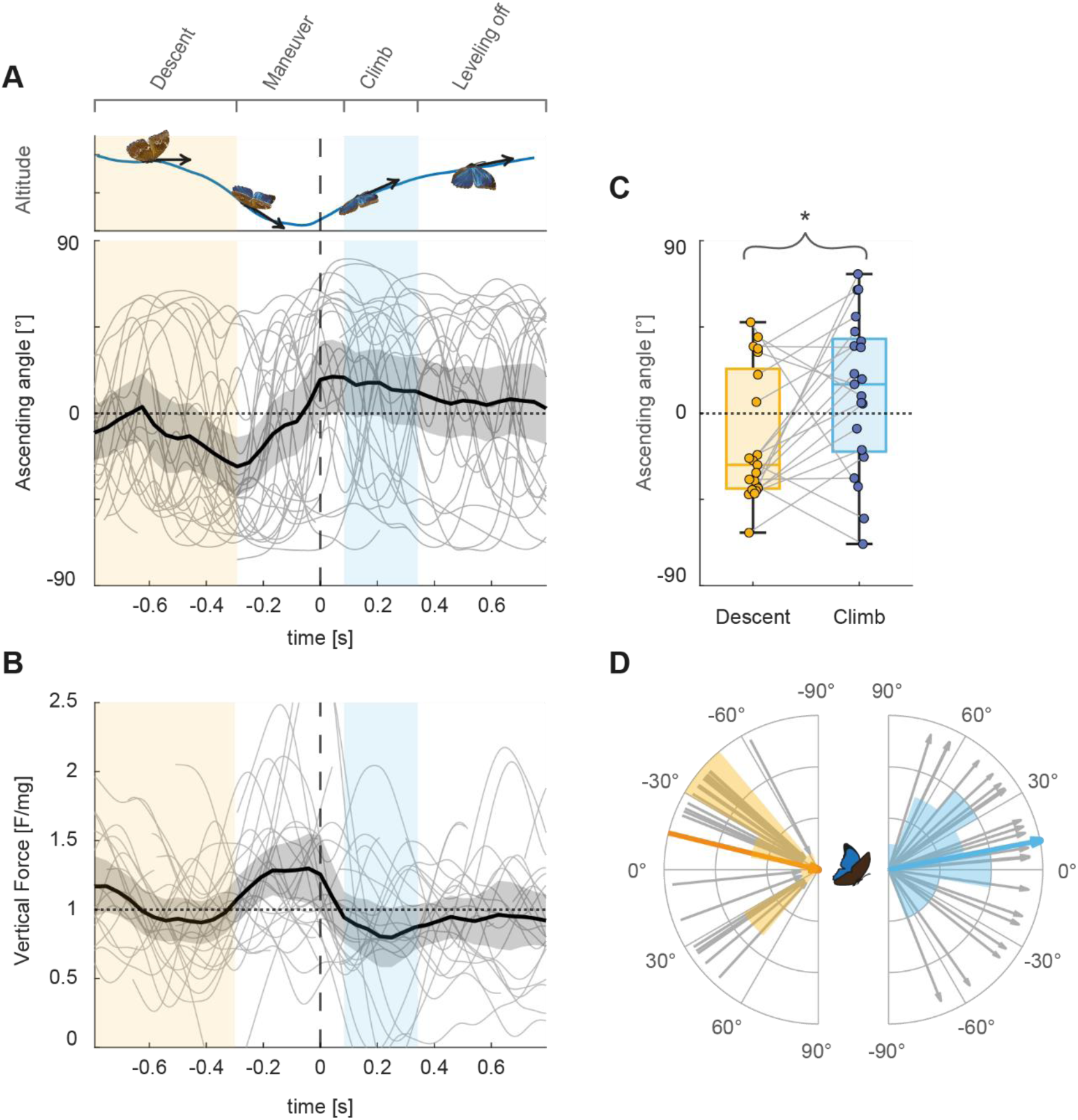
Vertical flight kinematics of 32 butterflies approaching a Morpho mimic and escaping from an insect net attack. Throughout the figure, we define four flight phases: the descent flight phase towards the morpho mimic (yellow), the rapid upward reorienting maneuver, the climbing flight following the maneuver (blue) which initiates escape, and the final phase during which the butterflies level off their flight for the rest of their escape. **A, B)** Temporal dynamics of ascending angles and vertical aerodynamic force before and after the attack (t=0). Data of individual butterflies are represented by thin grey curves; the thick black line with grey shading shows the mean and 95% confidence interval, respectively. The maneuvering phase spans from -0.3s to 0.1s. To illustrate the resulting theoretical flight that corresponds to the mean ascending angle curve, a vertical position curve has been generated on top. During the maneuver, vertical force production peaks at 1.3g resulting in a climbing maneuver; this is followed by a decrease in force production to <1g, resulting in a leveling off in climb angle. **C)** Boxplots of the ascend angles during the descent phase (orange) and the climb phase (blue), showing a significant mean ascend angle increase of 25° (p=0.016, ES=0.602, n=21). **D)** Polar representation of the mean ascend angles measured during the descent phase (yellow) and climb phase (blue). Individual mean ascend angles are represented by the thin grey arrows, while the overall mean angles are represented by the thicker colored arrows (orange: -13.8° during the descent phase; blue: 10.9° during the climb phase). Ascend angle distributions during both phases are shown by polar histograms.

Examining the variations of ascend angle throughout the flight sequences, we indeed detected a quick upwards angular shift prior to the moment of closest encounter to the net (*t*=0 s; fig. 2A). We tested whether this upwards shift was consistent across the filmed individuals by comparing the distributions of ascend angles during the descent and climb phases using a Wilcoxon signed-ranks test. This comparison shows a robust increase in climb angle of ∼25° (fig. 2C,D and fig. S1A; *p*=0.016, *ES*=0.602; descent phase: γ=-14°±34°; climb phase: γ=11°±40°). This result suggests that the immediate response to the looming attack was a climbing maneuver initiated at *t*∼-0.3s, resulting in an upward-directed escape flight. Adapting the same comparison to include the entire escape sequence (climb + leveling-off phases) yielded a non-significant difference (*p=*0.211, *ES=*0.304), suggesting that this upwards maneuver was only transient.

The temporal dynamics of the weight-normalized vertical force produced by the butterflies also revealed a distinct vertical force peak during the maneuver phase (fig.2B). During the majority of the approach and escape flight (i.e. climb + leveling off phases), the mean vertical force production remains constant at approximately 1 g for weight-support, resulting in negligible wingbeat-average vertical accelerations. In contrast, during the maneuver phase (*t*∼[-0.3; 0.1]), the mean vertical force production peaks at ∼1.3 g, which explains the rapid change in ascend angle, from a descent approach into a rapid climbing escape flight. Thus, butterflies only increased their vertical force production when coming into close contact with the looming net, but did not produce an increased vertical force for the rest of their escape.

The vertical escape dynamics was independent of both the direction of approach and attack, as shown by the lack of significant effect in the associated GLM (approach: *p*=0.146; attack: *p*=0.165; interaction: *p*=0.233).

### Heading kinematics: *Morpho menelaus* butterflies escape in random directions

Secondly, we focused on horizontal escape flight kinematics (fig. 3). The overlay of all flight tracks combined shows a high variation in escape flight headings, and rather tortuous flight tracks post-attack (fig. 3A).

**Figure 3:**
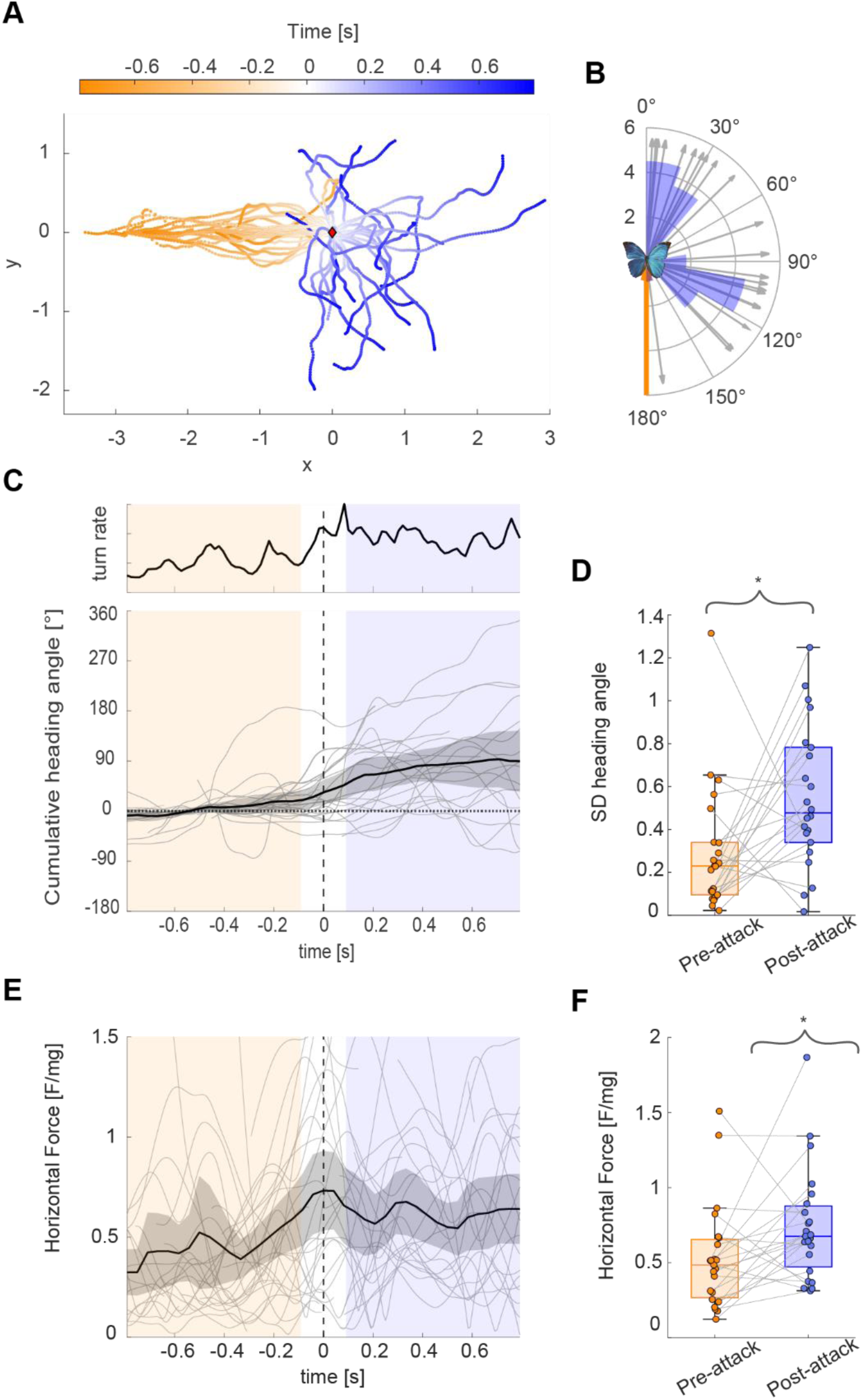
Horizontal flight kinematics of 32 butterflies approaching a Morpho mimic and escaping from a looming net attack. Throughout the figure, we define two flight phases: the pre-attack approach flight phase (*t*<-0.1 s; orange) and the post-attack escape flight (*t*>0.1 s; blue), separated by the evasive maneuver buffer zone. **A)** Top view of all flight trajectories, aligned at the moment of attack (*t*=0; red diamond). **B)** Polar representation of the evasive turn angle of each escaping butterfly, estimated as the mean heading angle during the first 30 video frames of the post-attack escape flight. The aligned pre-attack direction is represented by the thick inward orange arrow. Thin grey arrows show individual turn angles and proportions are illustrated by the blue polar histogram. The horizontal turn angles are highly variable, and clustered among either small turn angles (<40°) or large turn angles (>80°). The distribution is considered uniform around the semi-circle (V-test, *p*=0.120, ES=0.181, *n*=21). **C)** Temporal dynamics of turn rate (top) and heading angles (bottom) of the aligned and mirrored flight tracks. Thin grey lines show individual tracks; thick black lines and grey shading show the means and 95% confidence interval, respectively. Turn rate is defined as the time derivative of the average track heading. Turn rate peaks at just after *t*=0 s and then remains increased. **D)** Standard deviation of heading angles during both flight phases. The median variability in heading angles during the post-attack escape flight phase is 86% higher than during the pre-attack approach phase (*p*=0.024, ES=0.54, *n*=21). **E)** Temporal dynamics of the weight-normalized horizontal force production. Thin grey lines show individual tracks; thick black line with grey shading shows the mean and 95% confidence interval, respectively. Horizontal force peaks during the evasive maneuver (*t*∼0 s) and remains increased during the subsequent escape flight phase. **F)** Horizontal aerodynamic force production during both flight phases. The median weight-normalized force production during the post-attack escape flight phase (0.74g) is 40% higher than during the pre-attack approach phase (0.53g) (*p*=0.029, ES=0.52, *n*=21). For all boxplots, connected circles show paired data for each individual and boxplots show median and quartile ranges; stars show level of significant differences.

#### Evasive turn dynamics

To quantify the turning dynamics that underly the initial change in heading, we aligned all tracks, mirrored all left-hand turns (fig. 3B), and then determined the temporal dynamics of the turn rate, the cumulative changes in heading angle (fig. 3C), and the horizontal aerodynamic force production (fig. 3E).

This shows that the horizontal aerodynamic force peaks at *t*∼0 s, and the turn rate peaks at *t*∼0.1s, suggesting that the butterflies react to the attack not only by a climbing behavior, but also with a horizontal turn maneuver. The difference in timing of the horizontal and vertical force production suggests that the horizontal turn peaks approximately 0.1 s after the upward-directed response (figs 2C and 3C).

The distribution of mirrored turn angles (ϕ_mirrored_) is shown on figure 3B, and original turn angles (ϕ) in figure S1B. The absence of clustering around 0° (Fig. 3B; circular V-test: *p* = 0.120, ES = 0.181) indicates that butterflies generally perform a significant turn maneuver in response to the attack. Moreover, the distribution of turn angle magnitudes (ϕ_mirrored_) spanned almost the complete 180° range, as it varied from 4° up to 175° (fig. 3B). Among the distribution of ϕ_mirrored,_ two peaks stand out: one at relatively small turn angles and another at high turn angles. This suggests that escaping *Morpho menelaus* butterflies seemed to perform either small or large directional changes, with almost all turns being either lower than 40° or higher than 80°.

Furthermore, the Rayleigh test of uniformity applied on the original escape directions in the world reference frame (fig. S1C, ϕ) was non-significant (*p*=0.470, *ES*=0.175), suggesting no preferential escape directionality relative to the environmental context. Furthermore, neither the approach direction of the butterflies nor of the attacking net were found to influence escape direction, according to the corresponding GLM (approach: *p*=0.779; attack: *p*=0.209; interaction: *p*=0.653).

Overall, these results show that upon an attack, *Morpho menelaus* butterflies performed a rapid evasive turn maneuver in the horizontal plane, with variable turn magnitudes, and no preferential direction, suggesting a highly unpredictable escape direction.

#### Escape flight dynamics

During the post-attack escape flight phase, the intra-individual heading angle variability strongly increased as compared to prior the attack (fig. 3D; *p*=0.024, *ES*=0.54). Based on the mostly linear shape of the heading curves before the attack, the increase in heading variability suggests that the butterflies performed more and sharper turns during their escape than during their approach flight.

The post-attack flight is also characterized by a 40% rise in mean horizontal aerodynamic force production, consistently observed in all escaping butterflies (*p*=0.029, *ES*=0.52) (fig. 3E,F). The joint increase in horizontal force and heading variability suggests that increasing horizontal force production during the escape enables the butterflies to perform more and sharper directional changes, possibly enhancing flight path unpredictability.

### Horizontal flight path erraticity increases during the escape

To precisely characterize the unpredictability in the escape flight, we estimated the trajectory erraticity throughout both the approach and escape flight phases (fig. 4). We predicted an increase in erraticity during escape flight, enhancing escape by hindering the success of predatory follow-up attacks.

**Figure 4:**
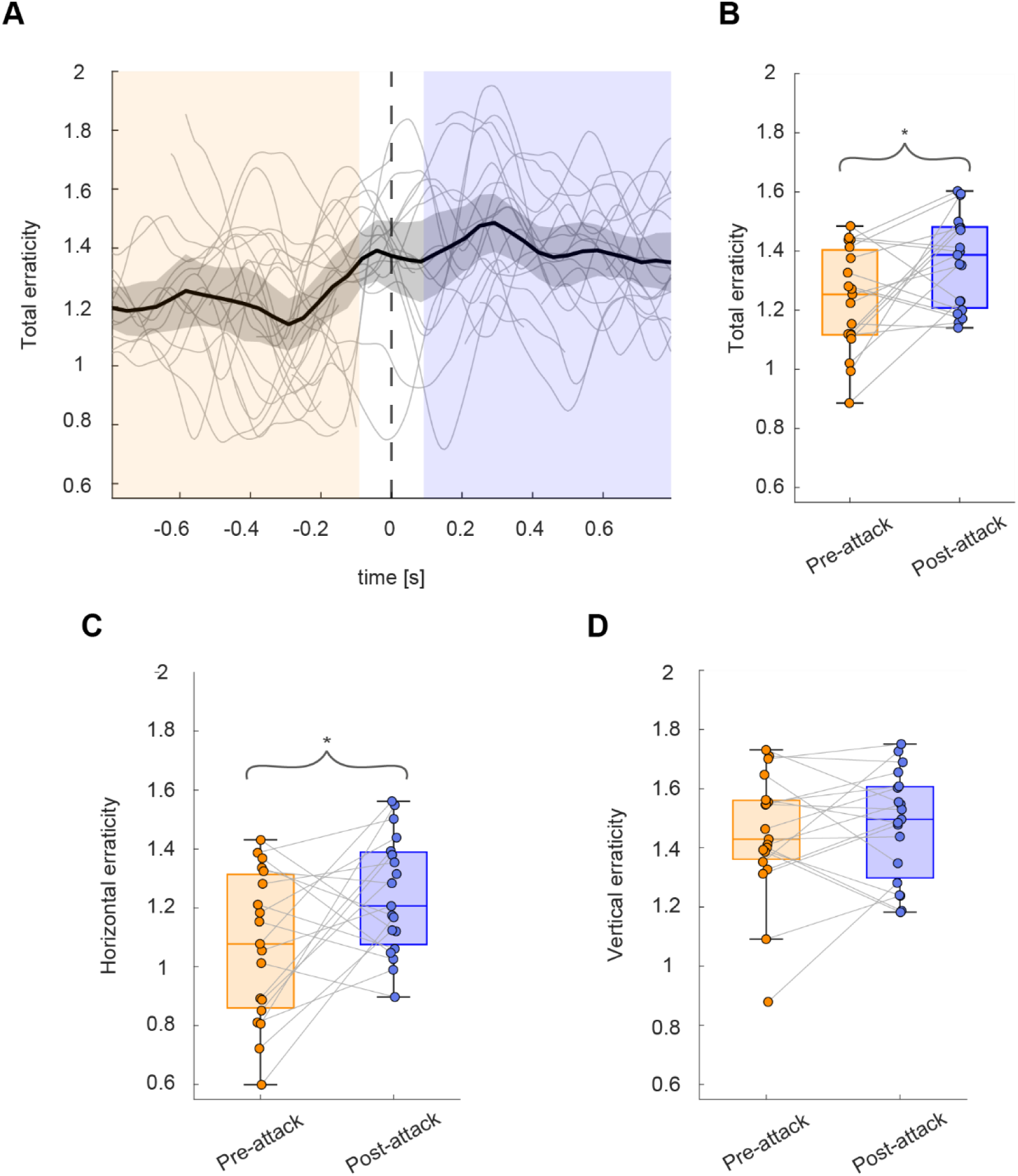
Flight path erraticity throughout the 32 approach and escape flights. Throughout the figure, we define two flight phases: the pre-attack approach flight phase (*t*<-0.1 s; orange) and the post-attack escape flight (*t*>0.1 s; blue), separated by the evasive maneuver buffer zone. **A)** Temporal dynamics of the total flight erraticity based on the three-dimensional flight trajectories. Thin grey lines show data for individual tracks; thick black line with grey shading shows the mean and 95% confidence interval, respectively. **B-D)** Total erraticity (B), horizontal erraticity (C) and vertical erraticity (D) during both flight phases. Connected circles show paired data for each individual and boxplots show median and quartile ranges; stars show level of significant differences. Both total and horizontal erraticity increase significantly from the pre-attack to the post-attack phase, while vertical erraticity does not (total erraticity: *p*=0.027, *ES*=0.58, *n*=21 (B); horizontal erraticity: *p*=0.049, *ES*=0.52, *n*=21 (C); vertical erraticity: *p*=0.66, *ES*=0.12, *n*=21 (D)).

The temporal dynamics of total erraticity based on the three-dimensional flight trajectories (fig. 4A) show a relatively low erraticity during the pre-attack approach flight phase (*t*<-0.1 s) followed by an increased erraticity during the subsequent post-attack escape phase (*t*>0.1 s). On average, the total erraticity in the escape phase was 11% higher than during the approach flight (fig. 4B; *p*=0.027, *ES*=0.58), suggesting that the escaping butterflies increased their flight path unpredictability during the escape.

We tested whether this increase in total escape erraticity was primarily achieved through horizontal or vertical displacements (fig. 4C, D and fig. S2F, I). Vertical erraticity remained rather stable throughout the complete flight (fig. S2I) and did not significantly differ between the approach and escape phases (fig. 4D; *p*=0.66, *ES*=0.116). In contrast, horizontal erraticity showed temporal dynamics very similar to the total erraticity (fig S2F): it rapidly increased during the evasive maneuver and remained high during the subsequent escape, with a 16% median increase between the two phases (fig. 4C; *p=*0.049, *ES*=0.52).

*Morpho menelaus* butterflies therefore increase their flight path erraticity during their escape flights, by performing more protean movements in the horizontal plane. These observations are consistent with the increase in both horizontal heading variability and horizontal aerodynamic force production (fig.3D,F). Overall, *Morpho menelaus* presents unpredictability in both its variable evasive turn dynamics and its erratic movements during the subsequent escaping flight.

### Horizontal flight speed decreases but force production increases during escape

We tested how flight speed and the net total aerodynamic force production varied throughout the flight sequences. We hypothesized that butterflies would increase their flight speed post-attack to fly away from the threat faster, increasing their escape success.

#### Flight speed variations

The temporal dynamics of speed show that butterflies approach the lure at a mean flight speed of 3.7 m s^-1^ (fig. 5A and fig. S2B). But surprisingly, speed decreases during the evasive turn maneuver, and then remains low throughout the subsequent escape flight phase, at a mean speed of 3.0 m s^-1^. This shows that, contrary to our initial hypothesis, *Morpho menelaus* flies slower when escaping a looming attack, with a significant average speed reduction of 19% (fig 5C; *U*_pre_=3.7±0.8 m s^-1^; *U*_post_=3.0±0.8 m s^-1^; *p*=0.031, *ES*=0.53).

**Figure 5:**
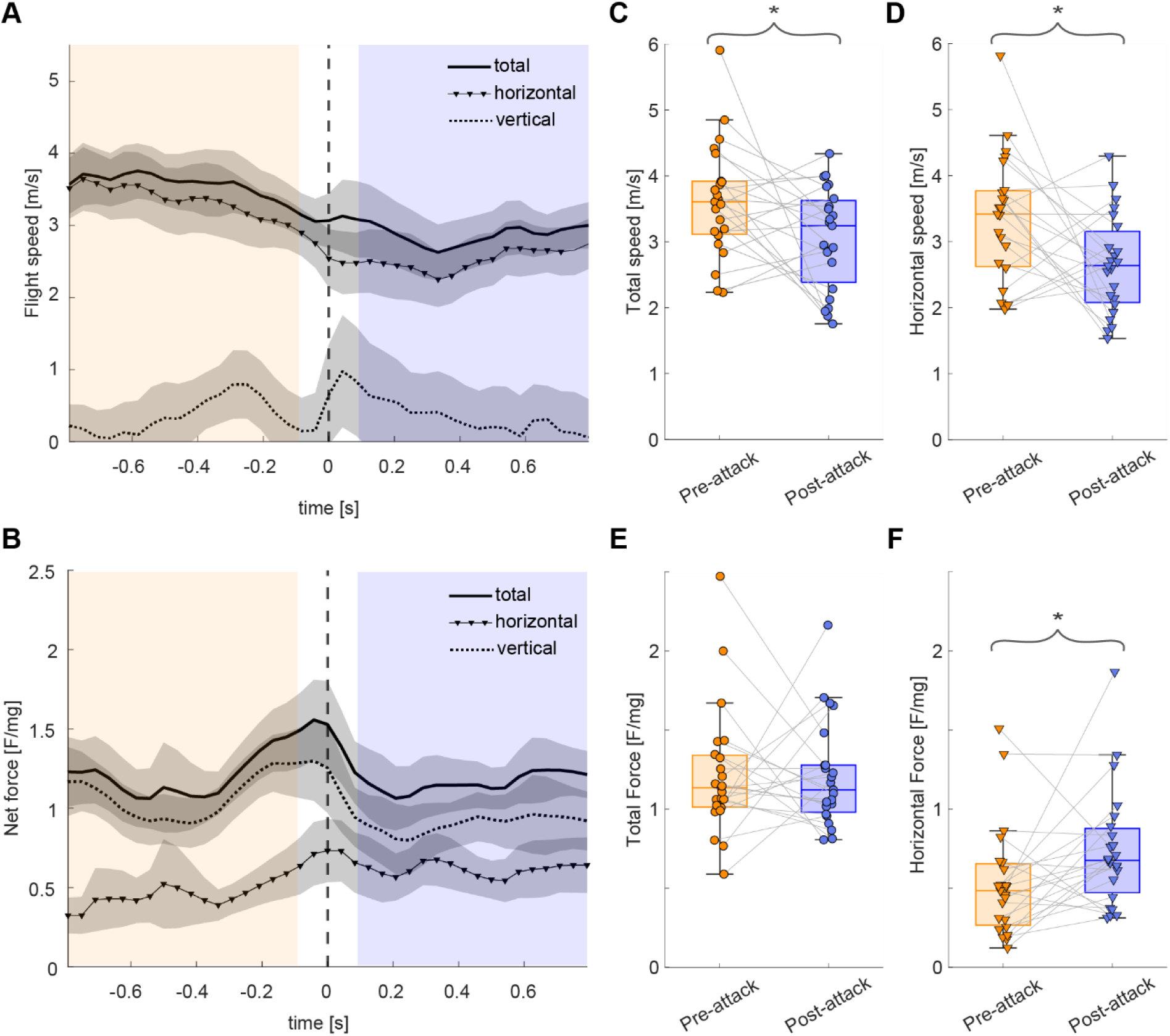
Flight speed and weight-normalized aerodynamic force production during all 32 recorded flights. Throughout the figure, we show total speed and force data using solid lines and circles, and the horizontal and vertical component with triangles and dotted lines, respectively. Moreover, we define two flight phases: the pre-attack approach flight phase (*t*<-0.1 s; orange) and the post-attack escape flight (*t*>0.1 s; blue), separated by the evasive maneuver buffer zone. **A,B)** Temporal dynamics of flight speed (A) and aerodynamic force production (B), whereby lines with grey shadings show mean data and 95% confidence intervals, respectively. **C,D,E,F)** Boxplots of mean total flight speed (B), mean total aerodynamic force (E) and their horizontal components (D and F, respectively), during the pre-attack and post-attack phases. Connected datapoints show paired data for each individual, and boxplots show median and quartile ranges; stars show level of significant differences. **C,D)** Both flight speed and its horizontal component decrease after the attack (total speed: *p*=0.031, *ES*=0.53, *n*=21 (C); horizontal speed: *p*=0.023, *ES*=0.54, *n*=21 (D)). **E,F)** The mean total force does not differ between the pre-attack and post-attack phases (p=0.76, ES=0.073, n=21 (E)); in contrast, the mean horizontal force production increases by 40% from pre-attack to post-attack (p=0.029, ES=0.52, n=21 (F)).

We then tested whether this decrease in escape flight speed was primarily achieved through a reduction in horizontal or vertical speed (fig. 5D and S2E, H). Vertical speed remained rather stable throughout the complete flight (fig. S2H) and did not significantly differ between the approach and escape phases (*p*=0.263, *ES*=0.27). In contrast, horizontal speed showed temporal dynamics very similar to the total speed (fig 5A and fig. S2E), as it decreased during the evasive maneuver and remained low during the escape. Total speed reduction thus mainly originates from a decrease in horizontal speed (fig. 5D; *p*=0.023, *ES*=0.54).

#### Aerodynamic force production

The variations in flight speed are driven by aerodynamic force production, and therefore we estimated the net weight-normalized aerodynamic force throughout all flight maneuvers (fig. 5B and fig. S2A). Total force remained rather constant throughout the pre-attack approach flight (*t*<-0.1 s) at a value slightly above 1g, peaked to approximately 1.5g during the evasive maneuver (−0.1<*t*<0.1 s), and then quickly dropped down again to ∼1g during the subsequent escape flight (*t*>0.1 s). As a result, the mean force production did not differ significantly between the pre-attack and post-attack flight phases (fig. 5E; *p*=0.76, *ES*=0.073).

We subsequently examined the temporal dynamics of the vertical and horizontal components of force (fig. 5F and fig. S2D, G). The peak and subsequent drop in total force detected during the evasive maneuver (t∼0 s) was primarily aligned with an equivalent peak and drop in vertical force production (fig. 5B), associated with the transient climbing response (fig. 2). The horizontal force component showed also a peak during the evasive maneuver, of approximately ∼0.7 g (fig. 5B). But in contrast with the total and vertical force, the horizontal force remained elevated after this evasive peak, resulting in a 40% increase in horizontal force production during the escape relative to the pre-attack phase (*p*=0.029, *ES*=0.52; figs 2E and 5F).

This increase in horizontal force does not show up in the total force as the normalized vertical forces decrease at the same time. As such, while the total force production does not increase in response to the attack, analyzing its subcomponents reveals that *Morpho menelaus* powers its escape flight by increasing its horizontal force production.

### Flight performance trade-off: erratic but slow escapes in *Morpho menelaus*

Our results show that the escape flights of *Morpho menelaus* have a reduced flight speed and increased flight path erraticity, both primarily due to variations in their horizontal components. At the same time, weight-normalized horizontal aerodynamics force production increased during the escape. This suggests that the increase in erraticity might be driven by this increase in aerodynamic force production. The decrease in speed might thus not result from a drop in force, but from a reallocation of force among movement dimensions (i.e. erratic turning instead of forward flight; fig. 6A).

**Figure 6:**
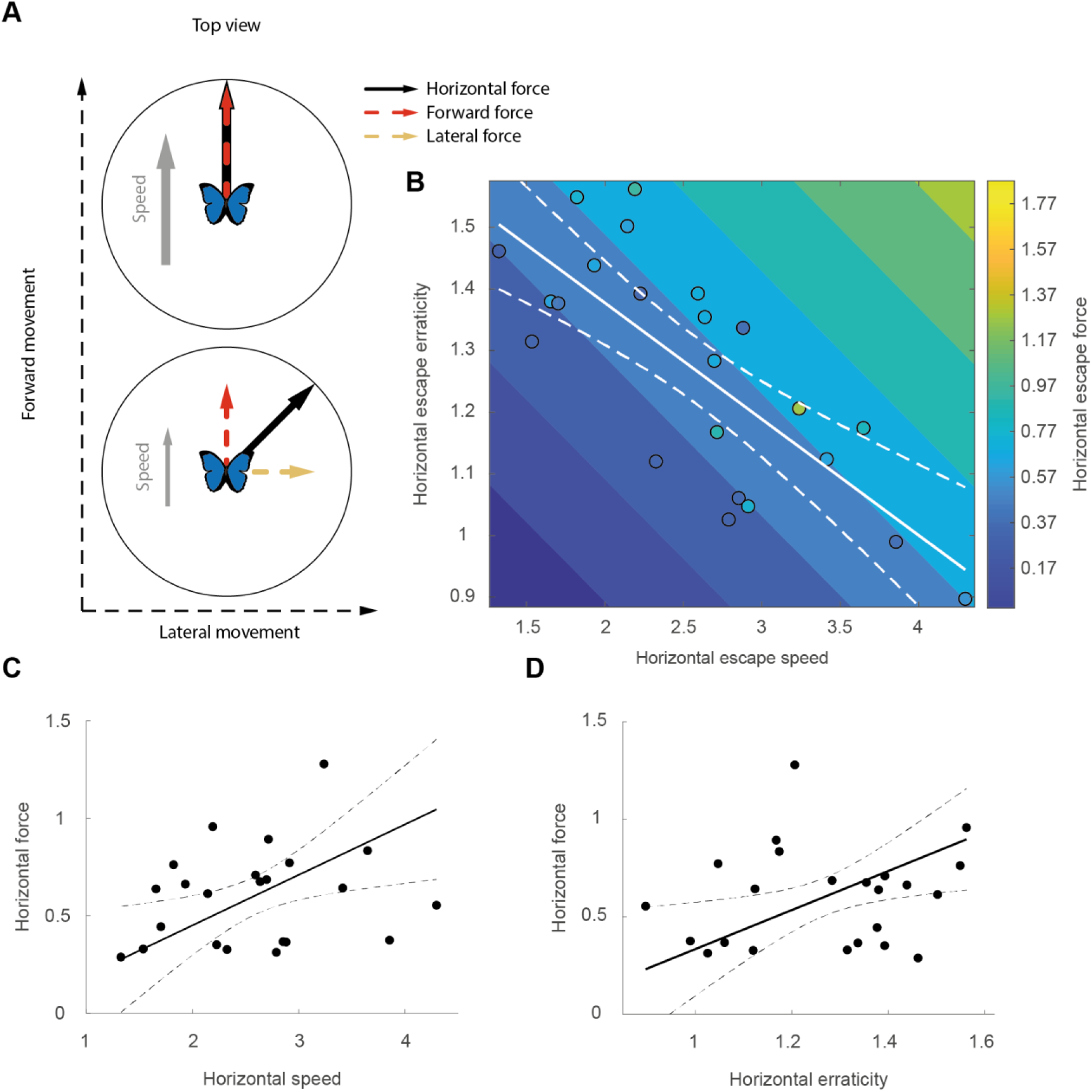
Escaping butterflies use horizontal force production to maximize erraticity and flight speed, leading to a trade-off between both components. Throughout, data points show results for individual butterflies; solid and dashed lines show predicted mean and 95% confidence interval of the different statistical models, respectively. **A)** Simplified free-body diagram representing the top view of a butterfly in flight. The horizontal force vector is represented by the thick black arrow and is separated into its forward and lateral components, illustrated by the red and beige dashed arrows, respectively. The resulting flight speed magnitude is represented by the grey arrow. The butterfly can increase its escape flight speed by aligning the force vector with the speed vector (i.e. producing forward thrust; top) or increase erraticity by producing oscillatory sideways forces (bottom). **B)** Combined results of the trade-off model (*E*_hor_∼*U*_hor_; white lines) and the combined force-dependency model (*E*_hor_+*U*_hor_∼*F/mg*_hor_; colored contour plot). The trade-off model (*E*_hor_∼*U*_hor_) shows a negative correlation between horizontal speed and erraticity, suggesting a trade-off between both aspects (*p*<0.001, ρ*_s_*=-0.72, *n* = 21). The combined force-dependency model shows a positive effect of horizontal force production on both speed and erraticity (speed: *p*=0.015, estimate=0.26; erraticity: *p*=0.019, estimate=1.003, *n* = 21). **C,D)** Linear model predictions for the separate force dependency models (*U*_hor_∼*F/mg*_hor_ at constant mean horizontal erraticity (B)); *E*_hor_∼*F/mg*_hor_ at constant mean horizontal speed (C)). According to these models, a 1g increase in horizontal force production can either increase flight speed with 3.86 m s^-1^ or increase erraticity with one unit.

This suggests a trade-off between flight speed and erraticity during the escape. We tested this hypothesis by first evaluating the correlation between flight speed and erraticity and secondly investigating the effect of force production on both parameters with a set of two GLMs (fig 6B). Here, we primarily focused on the horizontal components of flight speed, erraticity and aerodynamic force production, as these are the primary factors that vary during escape (see fig. S3 for the total and vertical components).

#### Trade-off between escape flight path erraticity and speed

First, we tested the correlation between the horizontal flight speed with horizontal erraticity during the escape flights. The results show that during escape, flight speed is tightly and negatively correlated with erraticity (fig. 6B; *p*<0.001, *r_s_ = -0.72*), whereby a 10% increase in erraticity results in a 15% decrease in flight speed. A highly similar relationship exists when we consider the total components of both flight speed and erraticity (fig. S3A; *p*<0.001, *r_s_=-0.76*), while no correlation was found for the vertical components (fig. S3C; *p*=0.56, *r_s_=-0.13*). This provides evidence of a trade-off between horizontal escape speed and flight path erraticity, but not for the vertical components.

Secondly, we tested the correlation between horizontal turn angle magnitude and flight speed before and after the attack. No correlation was found between turn angle magnitude and pre-attack speed (fig. S4A; *p*=0.42, *r_s_*=−0.18), implying that butterflies can perform turns of any magnitude regardless of their incoming flight speed. Similarly, no correlation was found with escape flight speed (fig. S4B; *p*=0.70, *r_s_*=−0.09). This suggests that the observed inter-individual variability in horizontal escape direction does not affect escape flight speed.

#### Horizontal force production drives escape flight erraticity

Next, we tested whether the observed increase in horizontal force production during the escape might produce the observed variations in flight speed and erraticity. To do so, we applied two generalized linear models to assess the relationship between horizontal force production and both horizontal speed and horizontal erraticity.

Despite the negative correlation between flight speed and erraticity, the horizontal force was positively related to both variables (speed: *p=*0.015, estimate=0.26, fig.6C; erraticity: *p*=0.019, estimate=1.003, fig.6D). Combining the results from both models into a single one (*E*_hor_*+U*_hor_*∼F*_hor_*/mg*; figure 6B) shows that *Morpho menelaus* produces, on average, horizontal force of approximately 0.7 g, that is used to produce a forward flight speed ranging from ∼1.5 to ∼4.5 m s^-1^, and a horizontal erraticity of ∼0.9 and ∼1.6. But individual butterflies operate within this domain along the diagonal (with *F*_hor_/*mg*∼0.7 g).

Such joint positive association between both erraticity and speed with force indicates a mechanistic biomechanical effect that underlies the speed-erraticity trade-off: escaping butterflies may use horizontal force to increase either their speed or path erraticity, via allocation of horizontal force to forward thrust or lateral oscillations, respectively (Fig 6A). Thus, when force production is limited, speed and erraticity compete for force resources, resulting in the observed trade-off. Hereby, *Morpho menelaus* seem to prioritize unpredictability over flight speed.

## Discussion

We characterized the escape flights of *Morpho menelaus* by recording the three-dimensional trajectories of 32 individuals responding to a looming attack in the wild. Escape responses combined a rapid upward maneuver and an unpredictable turn, followed by an erratic but relatively slow flight. Consistent with our first hypothesis, butterflies showed strong unpredictability in both direction and trajectory. Contrary to our second hypothesis, they did not increase speed but instead slowed down. This pattern reveals a trade-off between speed and erraticity, with *Morpho menelaus* favoring unpredictability during escape, supporting our third hypothesis.

### *Morpho menelaus* butterflies avoid attacks by flying upwards

In response to the simulated attack, *Morpho menelaus* generally first executed an evasive maneuver directed upwards relative to its initial flight direction. We interpret this rapid rise as an avoiding maneuver as it always preceded the moment of closest distance between the butterfly and the attacking net (i.e. *t*=0 s). More specifically, the maneuver initiation occurred ∼0.3 s before the moment of closest encounter, highlighting the rapidity of the butterflies’ response.

The onset of the net movement likely provided the butterflies with cues to anticipate the incoming attack. Such cues may be visual, due to the human attacker movement, the artificial nature of the lure once the butterflies approached close enough, the looming approach of the net (Peek & Card, 2016) or a combination of these. The airflow perturbations caused by the net’s movement may also participate in triggering the butterfllies’ reaction (Cribellier et al., 2022).

Other instances of upward escape in flying prey have been documented in birds and insects. Small birds with large flight muscles such as the common swift (*Apus apus*) commonly aim upwards to outclimb and escape heavier predators (Hedenström & Rosén, 2001). Dipteran insects, such as mosquitoes, have also been documented to escape traps with climbing flight (Cribellier et al., 2018). However, in *Morpho menelaus*, the upward direction was transient and did not persist throughout the escape. This suggests that the climbing maneuver may primarily avoid the initial strike rather than sustain escape. The upward-directed evasive maneuver may be a plastic behavioral adjustment to the attacker’s terrestrial position and the upward direction of the net. However, the lack of correlation between attack and escape direction does not support this hypothesis.

The climbing maneuver may thus be an instinctive response to deviate from the attacker’s path as quickly as possible, shaped by natural selection. The evolution of such a climbing reflex may be promoted in butterflies with large wings due to the upward acceleration generated by each downstroke (Fei & Yang, 2016). Insectivorous birds hunting Morpho butterflies (i.e. jacamars and kingbirds; Chai, 1986; Pinheiro, 1996) usually intercept flying insects from an elevated position. Because *Morpho menelaus* flies low in the understory, most attacks likely come from above. In response to such diving attacks, a climbing maneuver may be an efficient way to jink a predator during interception. Such behaviors have been referred to as a “turning gambit” (Corcoran & Conner, 2016; Howland, 1974), enhancing prey survival probability.

#### The horizontal escape turn dynamics of *M. menelaus* are unpredictable

Directly following the climbing maneuver, *Morpho menelaus* performed a horizontal evasive turn. These turns span a wide range, from almost 0° to 180°, with a clear separation between two groups: 9 out of 21 individuals performed minor turns of less than 40°, and 10 performed turns larger than 80°. We found no statistical evidence of a dependence of turn angle on initial heading, attack angle, or initial flight speed. Similarly, escape turn magnitude did not affect subsequent flight speed, suggesting that horizontal escape angle is not constrained by speed or by the direction of the incoming attack.

At the environmental scale, we also found no consistent patterns in escape direction. Given the experimental conditions, butterflies could escape along the open riverbed or toward the forest understory. The latter likely provides a safer environment, as dense vegetation limits bird attacks and favors camouflage. However, the lack of directional preference suggests that *Morpho* did not rely on sheltering within the forest (Wirsing et al., 2010).

Taken together, these results demonstrate the high unpredictability of horizontal escape direction in *Morpho menelaus*, limiting any directional bias that predators could learn and exploit (Jabłoński & McInerney, 2005).

Despite this apparent advantage, the framework proposed by Domenici et al. (2011a) suggests that completely random escape directionality is not always advantageous against an intercepting attack. Randomness relative to attack angle includes the possibility of fleeing toward the attacker, increasing capture risk. Species documented to perform fully random escapes also tend to lack fine motor control (e.g., mosquito larvae: Brackenbury, 1999; springtails (Collembolans): Christian, 1978), suggesting that complete randomness may reflect motor limitations rather than adaptation.

However, Domenici’s model is based on two-dimensional data, and a simple extension to 3D—where advantageous escape directions would form a hemisphere rather than a half circle directed away from the attacker—may be misleading (Møller, 2010). In 3D, energetic asymmetries between horizontal and vertical movement constrain both predator and prey performance (Hedenström & Alerstam, 1992). In the absence of a 3D null model, interpreting the observed unpredictability of horizontal escape directions in *Morpho menelaus* remains challenging. Nevertheless, unlike mosquito larvae and springtails, butterflies are highly maneuverable, suggesting that their escape randomness is not a consequence of poor motor control. Instead, this high directional unpredictability might bear some advantage against predation.

#### *Morpho menelaus* increase horizontal erraticity during their escape flight

Following the rapid climbing and turning maneuver, *Morpho menelaus* performed distinct escape flights away from the attack site. These flights have often been described as “erratic” based on subjective perception (Chai & Srygley, 1990; Jantzen & Eisner, 2008; Marden & Peng Chai, 1991; Srygley & Dudley, 1993). Here, we provide a quantitative measure of flight path erraticity and show that *Morpho menelaus* increases erraticity after being attacked, resulting in more unpredictable escape flight trajectories.

Flying insects engaging in erratic movements can reduce predation risk, as shown by decreased attack rates in dragonflies preying on fruit flies (Combes et al., 2013) and lower capture success in bats targeting moths (Ter Hofstede & Ratcliffe, 2016). The effectiveness of this behavior relies on the difficulty of maintaining a precise interception course against unpredictably moving prey. A recent study on interception trajectories in birds of prey suggests that species relying on high-speed attacks (e.g., falcons) perform poorly against highly maneuverable prey (Brighton & Taylor, 2019).

In *Morpho menelaus*, erraticity increased only along the horizontal plane. This may reflect the lower energetic cost and faster execution of horizontal turns compared to repeated climbs and dives. While vertical erraticity did not increase after attack, its baseline value remained higher than the horizontal component. This elevated vertical variability may result from the relatively low wingbeat frequency of *Morpho menelaus* (∼6 Hz; Le Roy, Amadori, et al., 2021), which produces oscillatory vertical motion during flight. Butterflies accelerate upward during the downstroke and downward during the upstroke (Suzuki & Yoshino, 2018). Whether this vertical erraticity is a by-product of flight mechanics (as in silkmoths, Aiello et al., 2021) or an active behavior enhancing baseline unpredictability remains unclear.

#### Escape flight erraticity increases at the expense of a reduced escape flight speed

In our experiment, the escape phase was associated with a consistent reduction in flight speed. This is surprising, as speed is often considered a key determinant of escape success (R. S. Wilson et al., 2020). However, Clemente & Wilson (2016) showed using simulated prey and human predators, that escape success does not necessarily increase with speed, when reduced speed enhances maneuverability. In *Morpho menelaus*, decreased escape speed was indeed associated with increased horizontal erraticity, indicating a trade-off between speed and flight path unpredictability. Such trade-offs have already been documented, mainly in terrestrial locomotion (Del Simone et al., 2025; Wynn et al., 2015) and more rarely in flying insects (e.g., bumblebees, Mountcastle et al., 2015). To our knowledge, this study provides the first evidence of such a trade-off in butterflies.

This trade-off can be explained from a biomechanical perspective. Flying animals must generate aerodynamic forces to accelerate or maneuver. In the horizontal plane, deviations from straight flight require force production. Our results are consistent with this, as horizontal force was positively associated with both speed and erraticity. The trade-off likely arises from how this force is allocated: butterflies can increase flight speed by aligning the force vector with their velocity vector or increase erraticity by producing lateral oscillatory forces (Fig. 6A).

Given the importance of speed in escape success (Clemente & Wilson, 2016), our results highlight the role of protean flight in *Morpho menelaus’s* defensive behavior. Increasing erraticity rather than speed may be more effective against predators that easily outspeed their prey (Moore & Biewener, 2015), as shown in lucerne moths facing bats (Nakano & Mason, 2018). Erraticity may therefore be favored over speed by predator-driven selection, particularly in fliers with low wing loading such as Lepidoptera in general and *Morpho* in particular.

#### A coevolution of escape behavior and large blue wings?

The erratic escape flight behavior of *Morpho menelaus* documented here might have co-evolved with wing coloration. It has indeed been suggested that the wing iridescence observed in many Morpho species could facilitate the escape of these butterflies: the blue flashes produced by the alternate exposure of the dorsal bright and ventral cryptic wing surfaces might confuse predators (Young, 1971; Debat et al., 2018). Virtual predation experiments using birds (Silvasti et al., 2024) and humans (Murali, 2018; Murali & Kodandaramaiah, 2020) as predators found that high wingbeat frequency but also erratic flashing, especially in large individuals, increased attack errors. The increased escape flight path erraticity in *Morpho menelaus* should enhance the confusing visual effects of the bright blue flashes and increase the difficulty of capture. Natural selection may thus have jointly affected the evolution of iridescence and escape flight behavior in *Morpho menelaus*. Expanding the analysis of evasive flight to other *Morpho* species, especially presenting various colorations, should allow for testing this hypothesis.

#### Stereoscopic videography in the wild captures relevant natural behaviors

In this study, we used stereoscopic videography to precisely characterize the natural escape flight of *Morpho menelaus* in the wild. Although field-based recordings present significant challenges, they provide access to behaviors that are difficult to capture reliably in controlled laboratory settings (Combes et al., 2012). Such challenges include a limited filming frame in an open space, as well as tracking on images with variable lighting conditions and visually noisy backgrounds. These constraints likely explain why only a handful of studies have applied stereoscopic videography in field conditions with wild animals (Fabian et al., 2024; Khandelwal & Hedrick, 2022; Le Roy, Roux, et al., 2021). Ongoing progress in deep-learning-based tracking tools is expected to help overcome these limitations and incite more research in the wild (e.g. SAM2, Ravi et al., 2025).

Despite these technical difficulties, field recordings were essential to capture the full expression of *Morpho menelaus*’ evasive behavior. In particular, they allowed us to quantify the variety of escape directions, the increase in erraticity during escape flights, and the resulting trade-off with flight speed. Such responses would most likely be constrained in captivity, where spatial limitations necessarily restrict movement and bias escape trajectories. Moreover, working in the wild allowed us to study individuals without prior handling or confinement, reducing potential stress-induced effects on behavior. Overall, our results highlight the value of studying free-flying animals in their natural environment to accurately assess complex behavioral responses.

## Supplementary material

**Figure S1:**
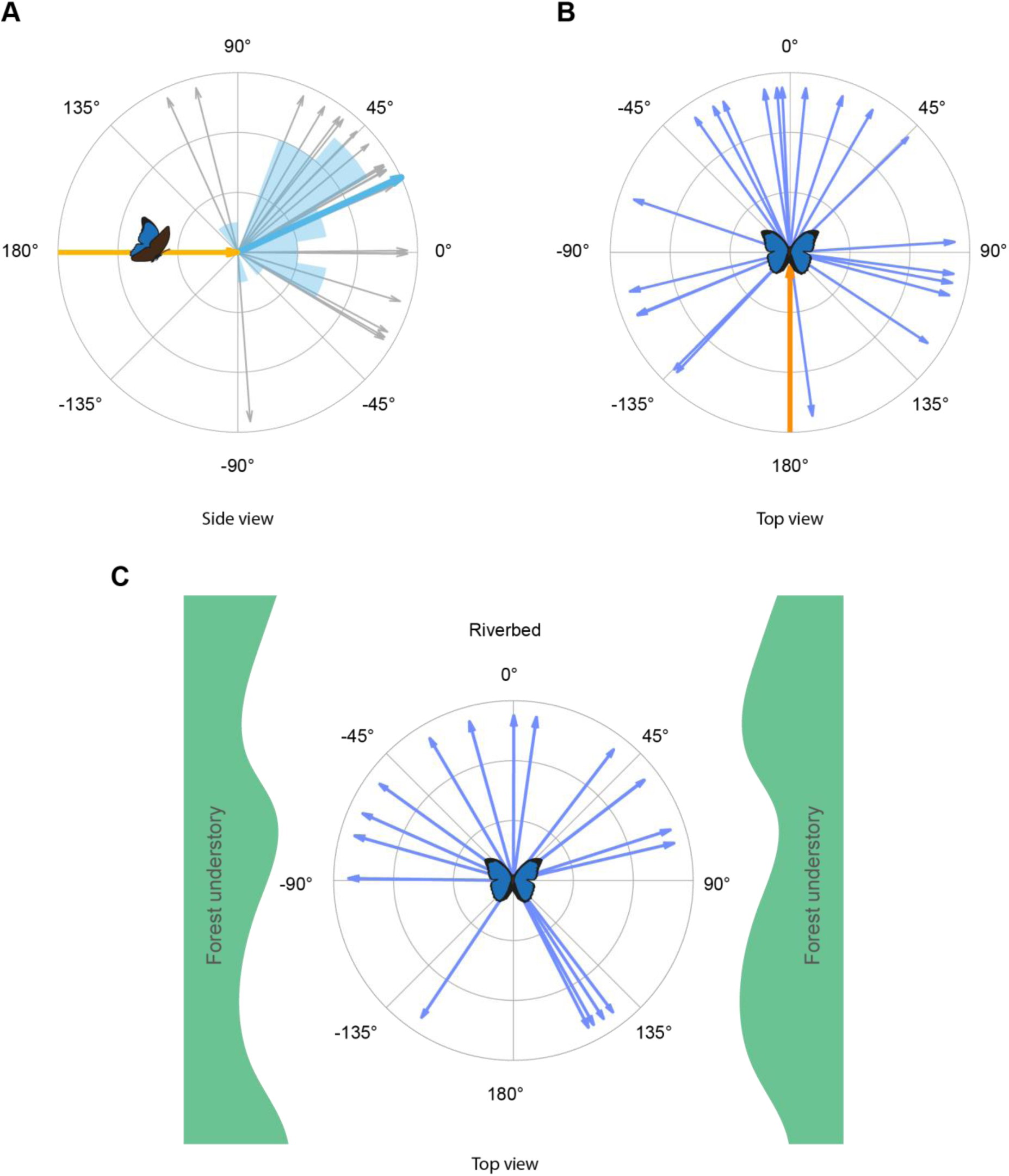
Escape directions of *Morpho menelaus* in the vertical and horizontal plane. All directions in **A** and **B** are relative to the approach angle before the simulated attack. **A)** Escape directions in the vertical plane, all grey arrows represent individual escape directions with the thicker blue arrow as the mean (25°). The yellow inward arrow represents the approach angle and is aligned at 0°. The mean blue arrow and the blue polar histogram illustrate the predominance of upward escape directions. **B)** Escape directions in the horizontal plane, all blue arrows represent individual escape directions relative to the approach flight (orange inward arrow) which are aligned at 0°. The variety of escape directions illustrates the lack of directional preference. **C)** Natural escape directions in the horizontal planes illustrating the local environmental context. The attack consistently took place in the middle of the riverbed which was aligned to the 0° orientation.

**Figure S2:**
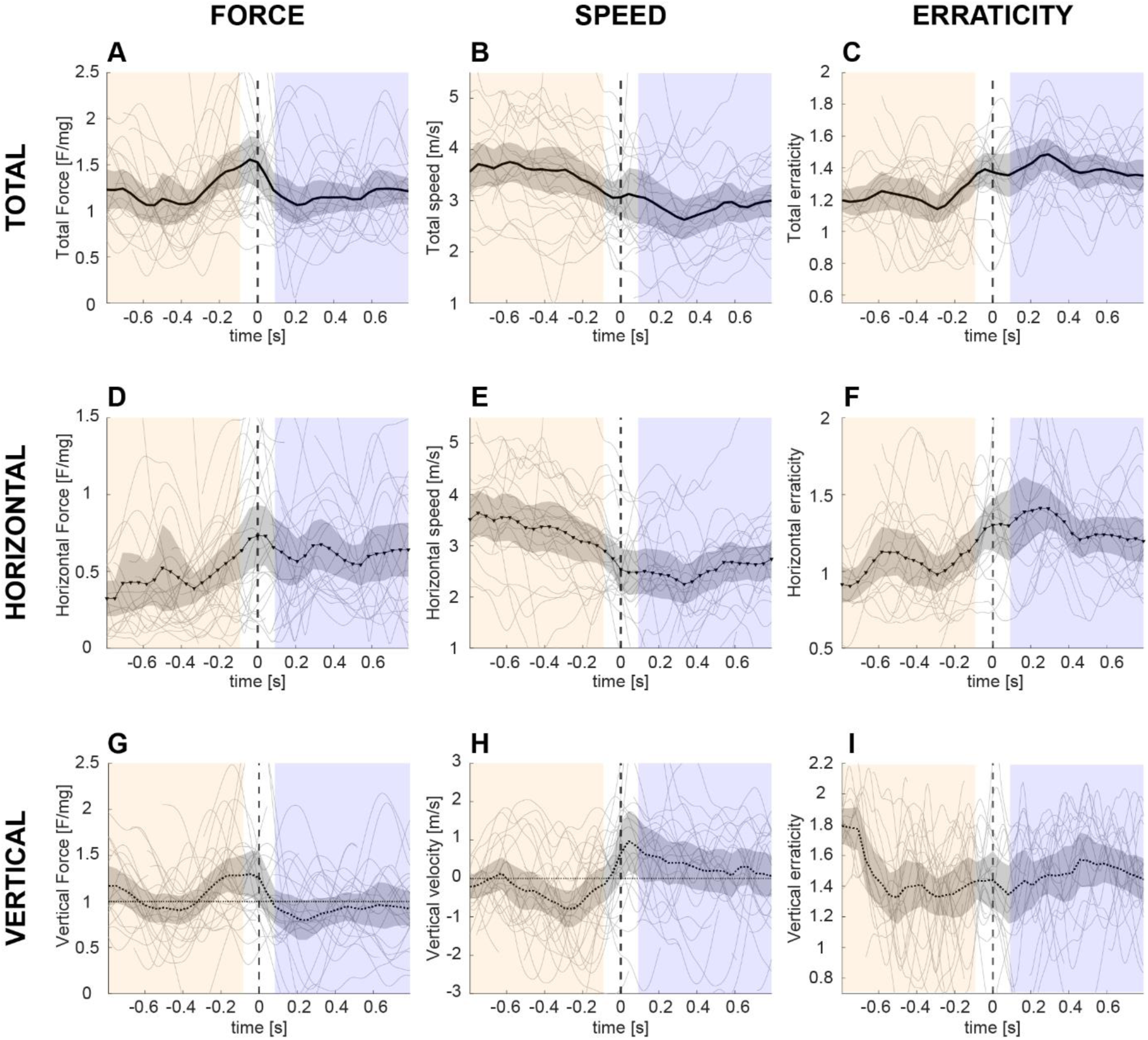
Temporal dynamics of speed, flight erraticity and force production of 32 *Morpho menelaus* butterflies. The temporal dynamics of all flight kinematics measured (speed, force production and erraticity) are represented along their total value (1^st^ row, **A**, **B**, **C**), their horizontal component (2^nd^ row, **D**, **E**, **F**) and their vertical component (3^rd^ row, **G**, **H**, **I**). For each graph, individual dynamics are represented by thin grey lines and their mean and 95% confidence interval by the thicker black line and grey shading. The mean line is solid for the total, punctuated with triangles for the horizontal component and dotted for the vertical component. The pre-attack and post-attack phases are illustrated by the orange and blue shadings respectively and separated by a 0.2s maneuver zone (t=[-0.1s : 0.1s]) with the moment of attack at t=0s.

**Figure S3:**
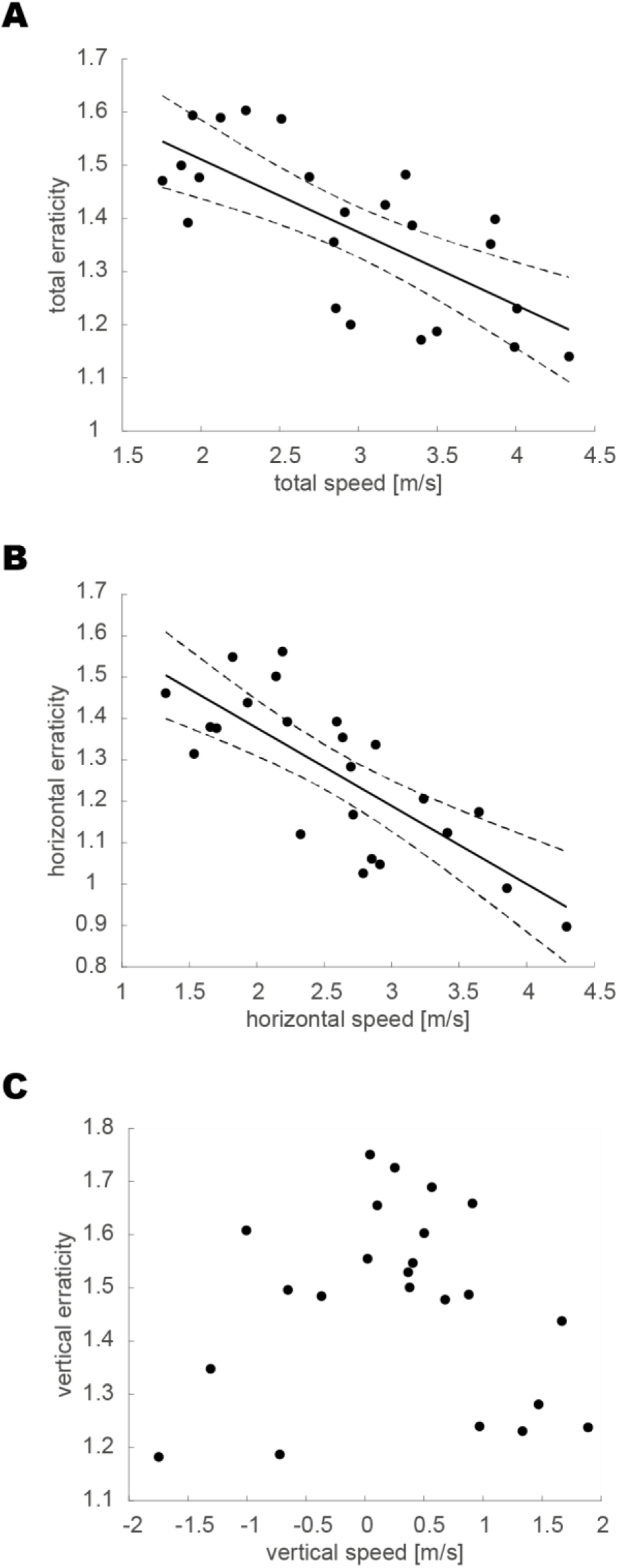
Results of the speed/erraticity trade-off models for the total, horizontal and vertical components. For each figure, individual butterflies are represented by the black points while the solid and dashed black lines represent the model’s prediction when the correlation is significant. This is the case for the total component (**A**, *p*<0.001, *r_s_*=-0.76) and the horizontal component (**B**, *p*<0.001, *r_s_*=-0.72) but not for the vertical component (**C**, *p*=0.56, *r_s_*=-0.13). These results indicate that flight speed and erraticity are engaged in a trade-off mainly driven by their horizontal components.

**Figure S4:**
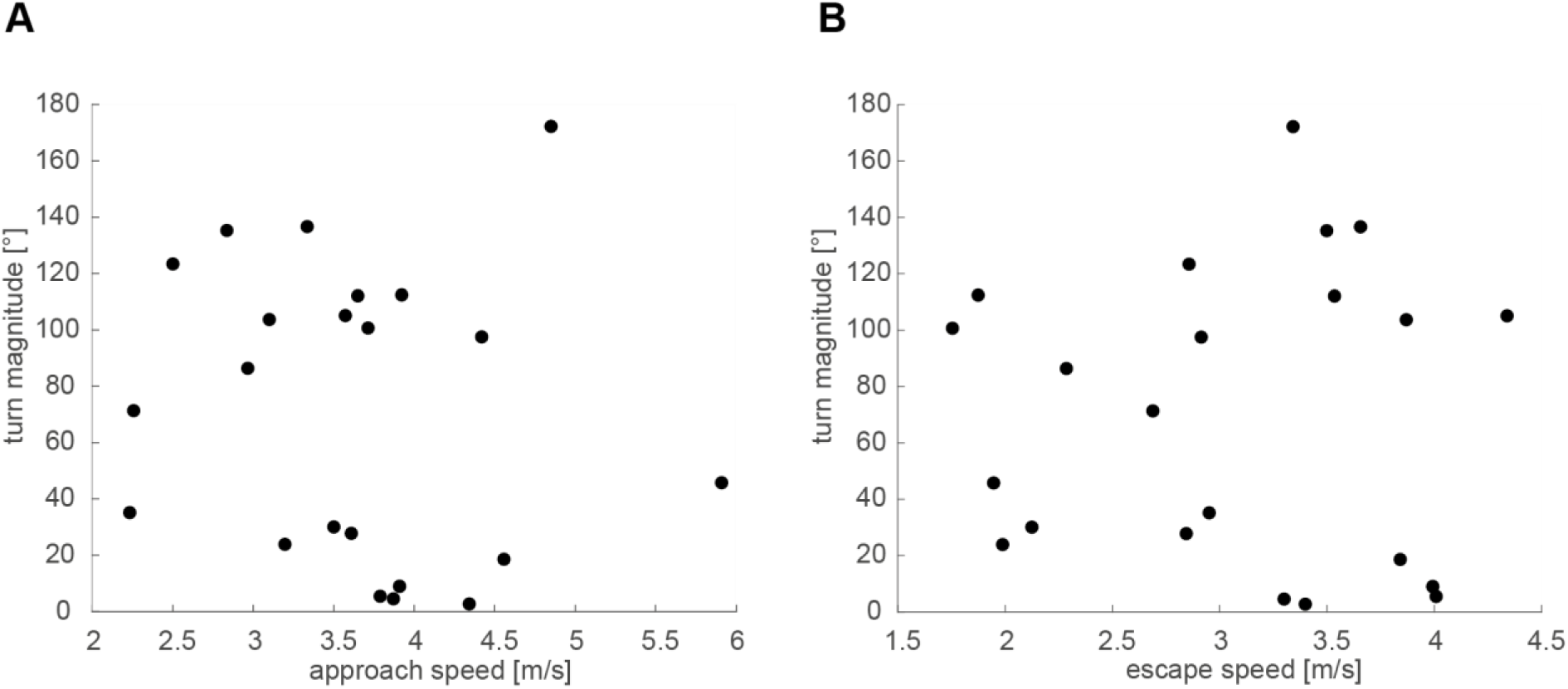
Relationship between turn magnitude during the evasive maneuver and flight speed before and after the attack. In both graphs, the black dots correspond to individual butterflies. Neither approach speed (**A**) nor escape speed (**B**) was significantly correlated to the evasive turn magnitude (**A**, *p=0.42,* r_s_*=−0.18*; **B**, *p=0.70,* r_s_*=−0.09*). These results first indicate that the sharpness of the turn engaged by the butterflies in response to the looming attack was not constrained by their approach speed. Secondly, the turn magnitude did not affect subsequent escape flight speed, indicating that there is no apparent trade-off between the redirection of the escape trajectory and escaping fast.

